# Palbociclib and fulvestrant act in synergy to modulate central carbon metabolism in breast cancer cells

**DOI:** 10.1101/348722

**Authors:** Benedikt Warth, Amelia Palermo, Nicholas J.W. Rattray, Nathan V Lee, Zhou Zhu, Linh T. Hoang, Anthony Mazurek, Stephen Dann, Todd VanArsdale, Valeria Fantin, David Shields, Gary Siuzdak, Caroline H. Johnson

## Abstract

Palbociclib, is a selective inhibitor of cyclin-dependent kinases 4 and 6 and used as a first-line treatment for patients with estrogen receptor positive breast cancer. It has been shown that patients have improved progression-free survival when treated in combination with fulvestrant, an estrogen receptor antagonist. However, the mechanisms for this survival advantage are not known. We sought to analyze metabolic and transcriptomic changes in MCF-7 adenocarcinoma breast cancer cells following single and combined treatments to determine if selective metabolic pathways are targeted during combination therapy. Our results showed that individually, the drugs caused metabolic disruption to the same metabolic pathways, however fulvestrant additionally attenuated the pentose phosphate pathway and the production of important coenzymes. A comprehensive effect was observed when the drugs were applied together, confirming the combinatory therapy′s synergism in the cell model. This study highlights the power of merging high-dimensional datasets to unravel mechanisms involved in cancer metabolism and therapy.

**Highlights:**

○ First study employing multi-omics to investigate combined therapy on breast cancer cells
○ Fulvestrant attenuates the pentose phosphate pathway and coenzyme production
○ Synergism of palbociclib and fulvestrant was confirmed *in vitro*
○ Altered key pathways have been identified

**eTOC Blurb:** Johnson et al. applied an innovative multi-omics approach to decipher metabolic pathways affected by single versus combination dosing of palbociclib and fulvestrant in estrogen receptor positive breast cancer. Key metabolites and genes were correlated within metabolic pathways and shown to be involved in the drugs′ synergism.

## Introduction

Cell cycle regulation is frequently disrupted in breast cancer (Cadoo et al., 2014; Mayer, 2015). Cyclin-dependent kinases (CDKs) control this regulation enabling quiescent cells the ability to enter the G_1_-phase and transition to the S phase. CDKs 4 and 6 phosphorylate the retinoblastoma (RB) protein enabling the release of E2F transcription factors (E2Fs) which mediates transition into the S-phase. Mutations to the CDK-RB1-E2F pathway typically result in the amplification of *CCND1* which encodes cyclin D1. Both are correlated with estrogen receptor positive (ER+) breast cancers. Thus, the re-establishment of normal cell cycle control through the inhibition of CDKs is an interesting option for the development of targeted cancer therapy. In recent years, agents have been developed which selectively target CDK4/6 (Mayer, 2015).

The development of CDK4/6 inhibitors such as palbociclib (Ibrance^®^, PD0332991) target the adenosine triphosphate (ATP) binding site of CDK4-cyclin D and CDK6-cyclin D complexes. This induces cell cycle arrest in the G_1_-phase (Johnson et al., 2016). Palbociclib is a selective, small-molecule inhibitor of CDK4/6 with the ability to block RB phosphorylation. It can be taken in combination with either the aromatase inhibitor letrozole, or the ER antagonist fulvestrant. Letrozole is used as initial endocrine-based therapy in postmenopausal women with ER+, human epidermal growth factor receptor 2 (HER2) negative metastatic breast cancer (Dhillon, 2015; Mechcatie, 2015). An accelerated U.S. food and drug administration (FDA) approval for this combined therapy was granted in February 2015. This was based on a randomized phase 2 study of 165 postmenopausal women which showed a progression-free survival (PFS) rate of about 20.2 months when treated with palbociclib and letrozole, compared to a PFS rate of 10.2 months among those treated with letrozole alone (Finn et al., 2015). In early 2016, the FDA approval for palbociclib was expanded to include combined therapy with fulvestrant, based on a phase 3 study (Turner et al., 2015). The median PFS of 3.8 months (placebo/fulvestrant) was increased to 9.2 months for palbociclib/fulvestrant (Turner et al., 2015). This combined treatment is currently in use for women with disease progression following hormonal therapy and can also be applied to pre-menopausal women (Cristofanilli et al., 2016).

*In vitro* studies have demonstrated the sensitivity of palbociclib towards different breast cancer cell lines and showed a synergistic effect with other drugs (tamoxifen and trastuzumab) in ER+ and HER2-amplified cell lines, respectively (Finn et al., 2009). In addition, a previous study investigated the effect of calcein acetoxymethyl-ester (a potent inhibitor of CDK4/6) on cancer cell metabolism, revealing an associated decrease in the concentration of sugar phosphates. This results in an imbalance of the pentose phosphate pathway (PPP) towards the non-oxidative branch versus the oxidative in human colon adenocarcinoma cells, and a state of metabolic inefficiency hypothesized to halt cell proliferation (Zanuy et al., 2012). We also recently employed metabolomics to investigate the effect of palbociclib and letrozole used in single and combination doses in MCF-7 cells. We determined that that the combined effects of palbociclib and letrozole on cellular metabolism had a more profound effect than each agent alone with enhanced changes seen to metabolites in nucleotide metabolism, amino acids, and central carbon metabolism (Warth et al., 2018).

Therefore, the aims of our study were to determine whether the combined effects of palbociclib and fulvestrant also exert a synergistic effect on breast cancer cell metabolism, and help understand the mechanism underlying the increase in progression-free survival for patients undergoing combined CDK4/6 inhibitor and endocrine therapies. We integrated metabolomic and transcriptomic data using XCMS Online to provide a multi-omic view of dysregulated metabolic pathways (Huan et al., 2017). This revealed the response of both metabolites and genes to each drug, and the combined effect of attenuating multiple pathways arresting cell growth.

## Results & discussion

### Metabolomics analysis

To identify metabolic pathways modulated by the drugs, breast cancer cells were dosed with either vehicle (control), palbociclib, fulvestrant, or a combination dose containing both drugs (Figure 1). Cells were harvested two and seven days post-dose, with another set harvested after seven days post-dose with refeeding on day four. This design allowed for the evaluation of short-and long-term responses to drug treatment. Untargeted mass spectrometry-based metabolomics was carried out on the cell lysates to analyze intracellular metabolites and the obtained data evaluated using the XCMS Online platform https://xcmsonline.scripps.edu (Forsberg et al., 2018; Gowda et al., 2014; Huan et al., 2017). The data revealed that metabolites in control samples (vehicle) changed in abundance over time due to expected cancer cell proliferation (Figure 2). Therefore, meta-analyses were used for further data evaluation, comparing altered metabolites from each drug treatment normalized to controls (Benton et al., 2008). The putatively identified metabolites from those experiments were subsequently mapped onto metabolic pathways using mummichog analysis housed on XCMS Online (Huan et al., 2017; Li et al., 2013). Metabolite identities were confirmed by comparison of MS/MS spectra with authentic reference standards, and quantified by targeted multiple reaction monitoring mass spectrometry, which provided additional validation.

**Figure 1.**
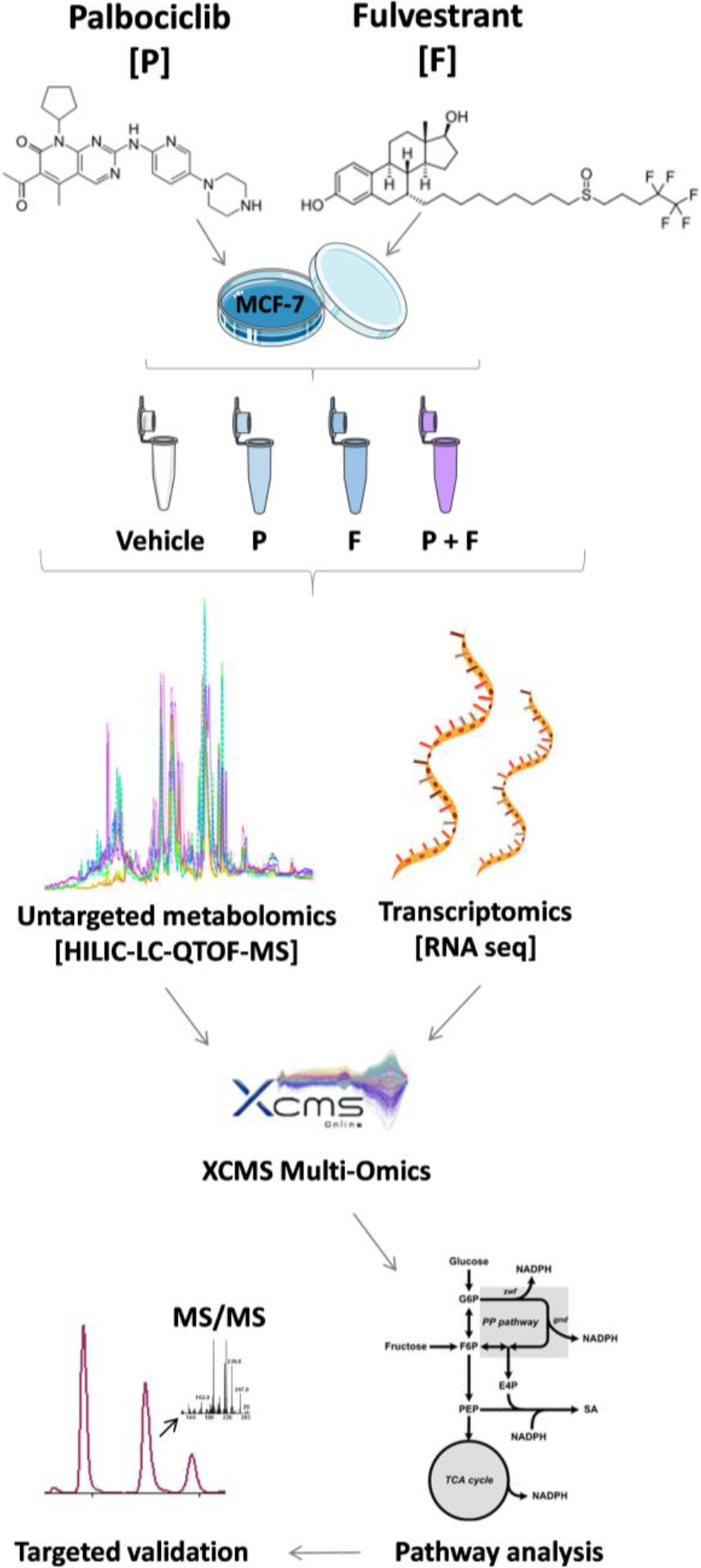
Overview of the workflow applied. MCF-7 breast cancer cells were treated either with the CDK4/6 inhibitor palbociclib, the estrogen receptor antagonist fulvestrant, or a combination of both drugs. Cells were extracted and analyzed after two days, seven days (without refeed), and seven days (with refeed).

**Figure 2.**
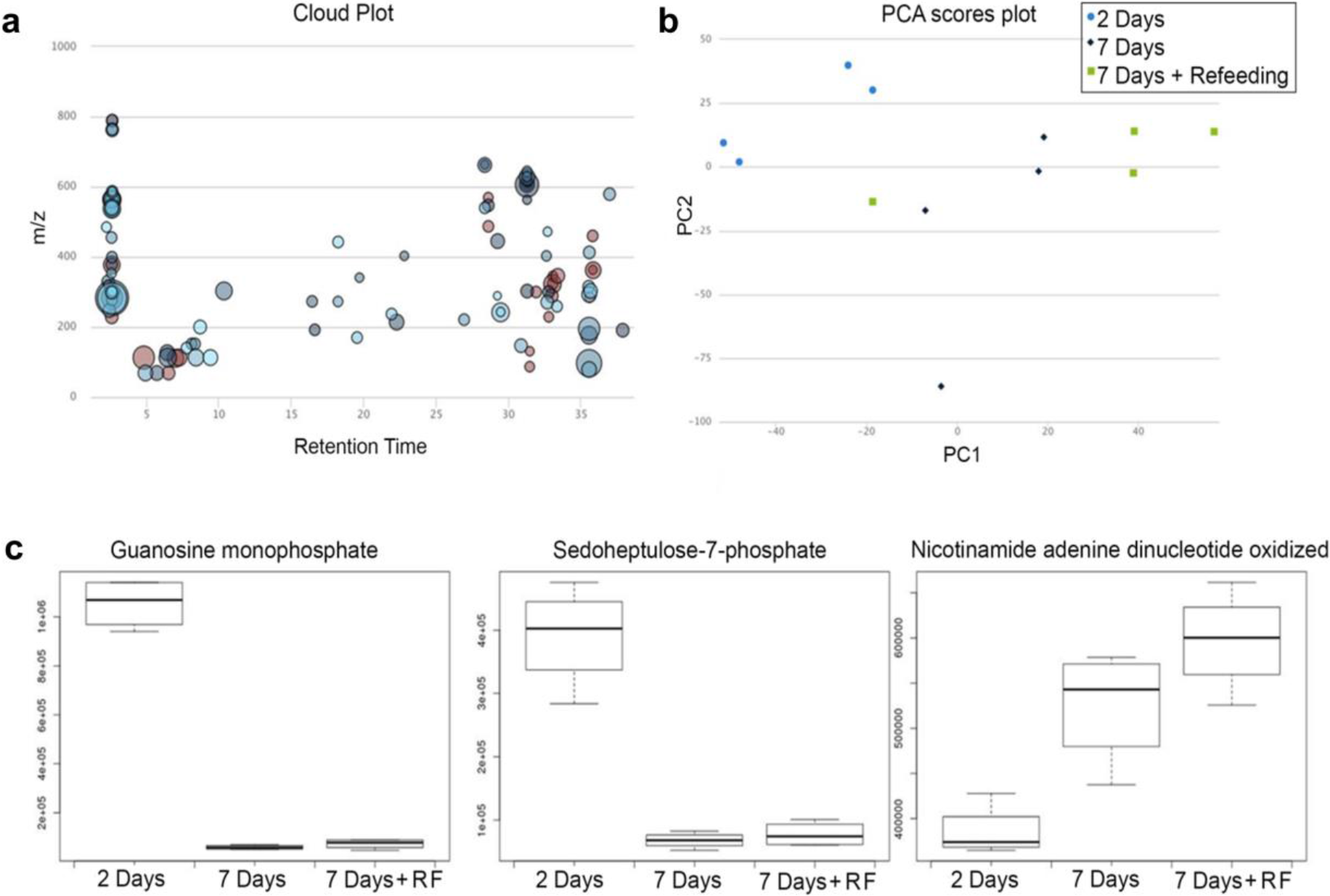
Multi-group analysis showing changes of the metabolome in MCF-7 cells treated with the vehicle (control). Cells were analyzed at two days, seven days and seven days with refeeding (n=4/group). Figure shows a) the cloud plot of all dysregulated metabolites compared across the different time points (fold change > 1.5, p < 0.01, fold change is represented by the radius of each feature in the plot), b) PCA scores plot, and c) box plots showing guanosine monophosphate, sedoheptulose-7-phosphate and nicotinamide adenine dinucleotide oxidized abundances change over time.

The results indicated that two days after a simultaneous dose of both drugs (palbociclib and fulvestrant) metabolites were dysregulated in multiple metabolic pathways in central carbon metabolism: the tricarboxylic acid (TCA) cycle, PPP, purine synthesis, and glycolysis (Figure 3). In the TCA cycle, succinate was increased 1.9-fold, and fumarate and malate were decreased 2.0-fold and 34.0-fold respectively. This indicates a blockade of succinate metabolism possibly through inhibition of succinate dehydrogenase (SDH). This effect was also evident in single agent dosing of both drugs, but malate was only decreased 4.0 and 3.5-fold when treated with palbociclib and fulvestrant alone, while combination dosing had more pronounced effect on malate depletion. Metabolites in both the oxidative and non-oxidative phases of the PPP were downregulated by combination dosing; sedoheptulose-7-phosphate (3.7-fold) and 6-phosphogluconate (2.5-fold). These metabolites were also downregulated by fulvestrant dosing alone to a similar extent but were unaffected by palbociclib, therefore it appears that the combination drug dosing effects on the PPP are driven by fulvestrant alone. Furthermore, three additional metabolites with roles in central carbon metabolism, were altered after combination dosing but were not changed with any single agent dosing; N-acetylglucosamine phosphate (1.2-fold increase), inosine monophosphate (16-fold decrease) and fructose-1-phosphate (2.8-fold decrease) (Table S1). Given the decrease observed in PPP, purine synthesis, glycolysis and TCA cycle intermediates after two days of dosing, these drugs act in synergy to target the same pathways that are important to cell growth and survival, as well as having separate actions on distinct pathways. This combinatorial effect enables a wider network of pathways to be modulated thus preventing the production of macromolecules and energy required for cancer cell growth, and increasing senescence. A list of all significant metabolite changes after two days is illustrated in Table 1.

**Figure 3.**
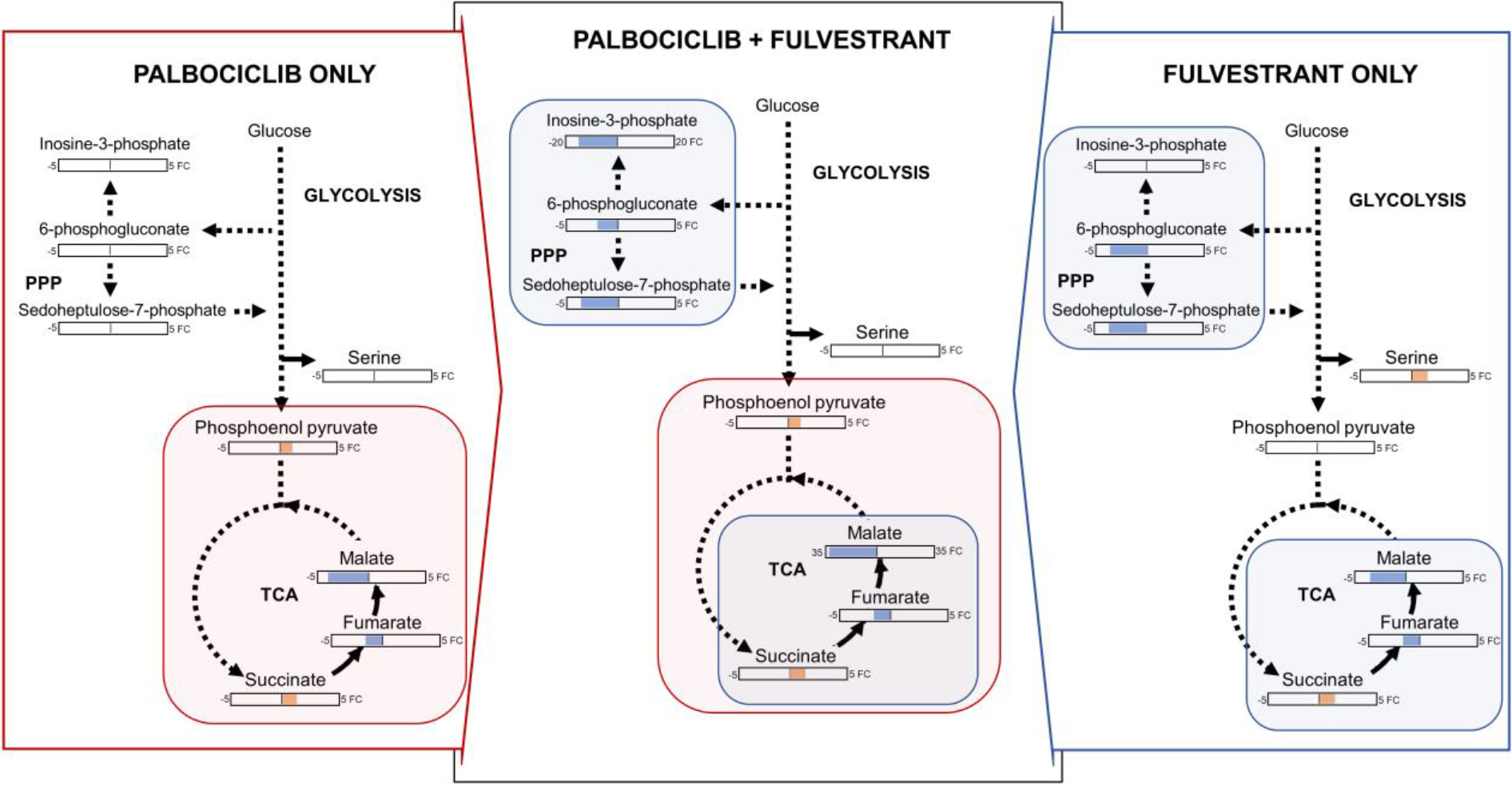
Overview of metabolite changes occurring in MCF-7 breast cancer cells two days post dosing with palbociclib, fulvestrant or in combination. Fold change values for significantly dysregulated metabolites (p < 0.05) are reported in orange (increase) and blue (decrease).

**Table 1.** Panel of significantly altered metabolites 2 days after dosing with palbociclib (n=4), fulvestrant (n=4) or a combination dose of both palbociclib and fulvestrant (n=4) when comparing dosed groups to control. Values are log transformed fold changes after quantification by multiple reaction monitoring, unpaired two-tailed t-test.

| Metabolite name | Palbociclib |  | Fulvestrant |  | Combination |  |
| --- | --- | --- | --- | --- | --- | --- |
|  | Fold Change | P-value | Fold Change | P-value | Fold Change | P-value |
| Succinate | 1.6 | 0.0025 | 1.8 | 0.0441 | 1.9 | 0.0021 |
| Malate | -4.0 | 0.0170 | -3.5 | 0.0381 | -34.0 | 0.0002 |
| Fumarate | -2.1 | 0.0189 | -2.1 | 0.0438 | -2.0 | 0.0003 |
| Phosphoenolpyruvate | 1.1 | 0.0450 | N.S | N.S | 1.2 | 0.0014 |
| 6-phosphogluconate | N.S | N.S | -3.7 | 0.0255 | -2.5 | 0.0496 |
| Sedoheptulose-7-phosphate | N.S | N.S | -3.6 | 0.0335 | -3.7 | 0.0003 |
| Serine | N.S | N.S | 1.7 | <0.0001 | N.S | N.S |
| Cholesterol sulfate | N.S | N.S | -5.2 | 0.0332 | N.S | N.S |
| Taurine | N.S | N.S | -2.1 | 0.0380 | N.S | N.S |
| Inosine monophosphate | N.S | N.S | N.S | N.S | -16.1 | 0.0052 |
| Fructose-1-phosphate | N.S | N.S | N.S | N.S | -2.8 | 0.0172 |

N.S. = not significant. Negative values indicate decreased fold changes

At seven days post-dose, changes to the metabolome were even more widespread than after two days. However, we first examined the effect of refeeding the cells at day four to determine if the observed changes were due to nutrient depletion. It was seen that after refeeding three of the metabolites were no longer dysregulated (malate, arginine, inosine monophosphate), thus they were not altered because of drug efficacy and increased senescence (Figure 4). There were however a number of metabolites that were changed after seven days of dosing and not affected by refeeding, as can be seen on Table S1. In contrast to the metabolic changes observed at day two post-dose, some of the metabolite concentrations were now changed in the opposite direction. For example, succinate decreased with both fulvestrant alone and combination dosing, however fumarate was not changed. This could be a result of decreased citrate/isocitrate utilization in the TCA cycle due to depleted nicotinamide adenine dinucleotide (NAD)+; of note citrate/isocitrate were increased under all dosing conditions at seven days. An additional change from two to seven days post-dose was seen in metabolites housed in the PPP; the intermediate sedoheptulose-7-phosphate was increased after fulvestrant and combination dosing. Fructose-1-phosphate was similarly changed, this metabolite feeds into the glycolytic pathway to produce glyceraldehyde and dihydroxyacetone phosphate. The change seen between day two and seven post-dose indicates the initiation of a salvage mechanism in which the cancer cells shift from glycolysis to PPP to increase production of nucleotides for energy, and glycerate-3-phosphate for serine biosynthesis, which were also decreased. In addition, the depletion of serine (fulvestrant and combination) and tyrosine (palbociclib, fulvestrant, and combination), and increase in phosphoenolpyruvate (palbociclib only) indicate that isoform M2 of pyruvate kinase (PKM2) regulation could be affected by the actions of palbociclib in single and combination doses (Amelio et al., 2014). Flux into the TCA cycle changed dramatically at seven days with an accompanying decrease in amino acids (aspartate, serine, proline, asparagine, tryptophan, tyrosine), NAD+ and nicotinamide adenine dinucleotide phosphate (NADP)+. All these actions were driven by fulvestrant and could lead to increased oxidative stress in the cell thus causing cell death (Vander Heiden et al., 2009).

**Figure 4.**
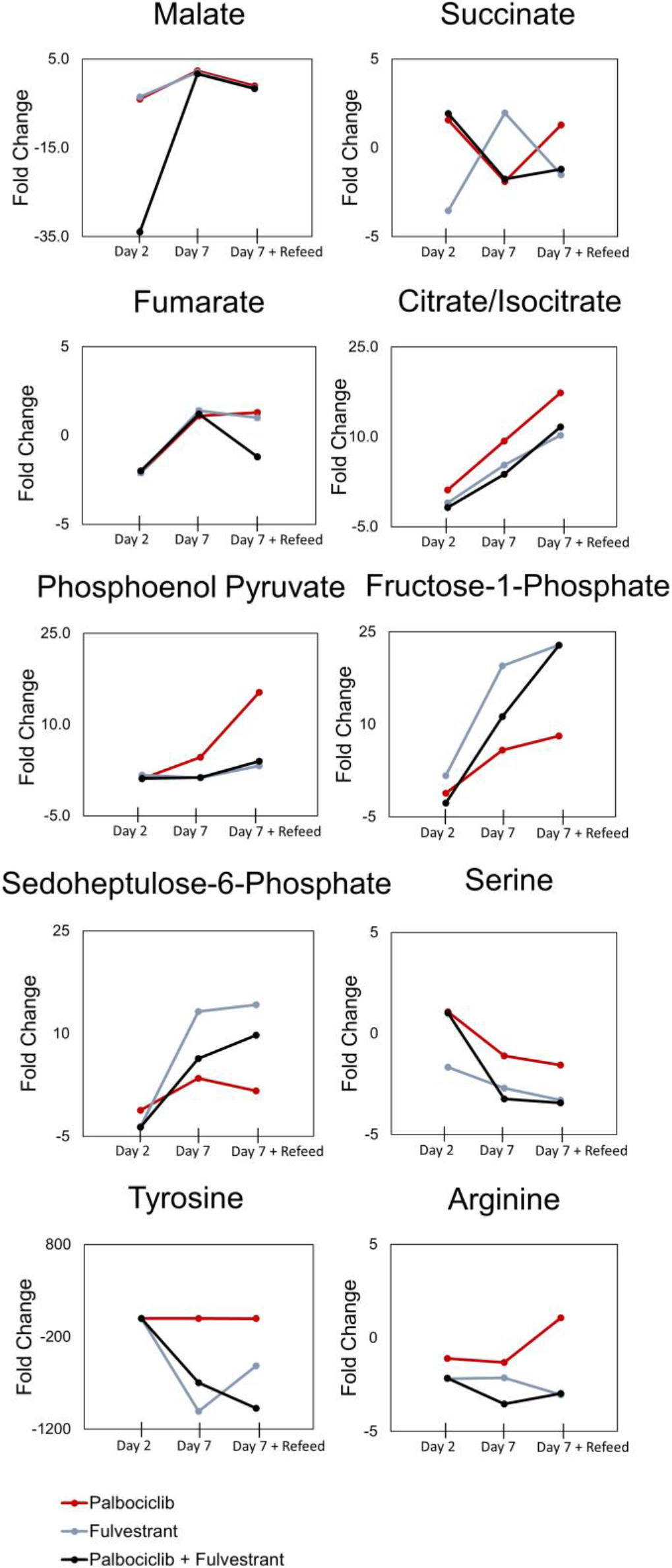
Overview of metabolite changes occurring in MCF-7 breast cancer cells with palbociclib and fulvestrant, single and combination dosing, after four days, seven days, and seven days with refeeding.

Taken together the results at seven days after combination dosing of palbociclib and fulvestrant show that the drugs act to inhibit purine, amino acid and important coenzyme synthesis. They also modulate glycolytic, TCA cycle and PPP metabolism possibly affecting the regulation of key enzymes such as PKM2 and SDH. In addition, changes to sedoheptulose-7-phosphate and fructose-1-phosphate levels from two to seven days post-dose indicate an adaptive mechanism diverting metabolic flux from glycolysis towards the PPP for the synthesis of important macromolecules and energy production (Vander Heiden, 2011). An overview of metabolite changes after seven days with combined dosing is shown in Figure 5.

**Figure 5.**
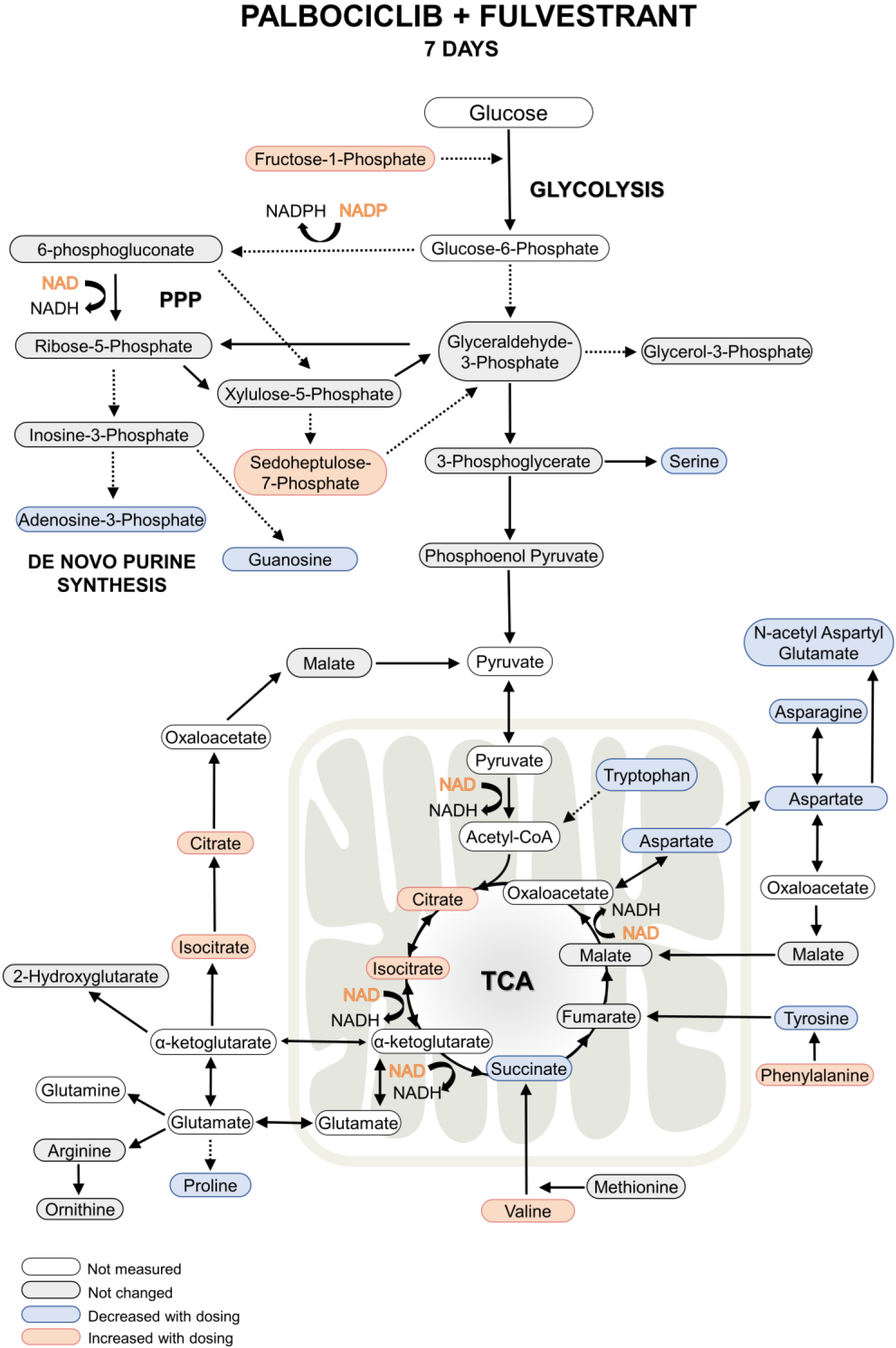
Overview of metabolite changes occurring in MCF-7 breast cancer cells. Seven days post palbociclib and fulvestrant, single and combination dosing.

### Transcriptomics analysis

In addition to comprehensive metabolomic analysis, RNA sequencing was performed to determine gene expression changes relating to single agent or combination drug dosing. Cells were collected at various time points (one day and seven days after dosing, with an additional time point at ten days with re-dosing at day seven). Metabolic genes appear highly over-represented among those whose expression were modulated by palbociclib as well as its combination with fulvestrant (Figure S1; Table S2). Gene set enrichment analysis (Kanehisa et al., 2012; Ogata et al., 1999) further identified significantly impacted metabolic pathways (Table S3) including metabolism of drug, nucleotides (both pyrimidine and purine; Figure S2), amino acids (histidine, phenylalanine and glycine/serine/threonine), fatty acids and retinol.

The differential regulation of selected genes after palbociclib, fulvestrant or combination treatment dosing compared to control can be seen on Figure 6 and Table S2. To correlate our metabolite changes with gene expression changes, we used the recently developed systems biology, omic data integration tool on XCMS Online (Huan et al. 2017). This tool identified four metabolic pathways which had both metabolite and gene involvement during combination dosing, these pathways are all associated with purine biosynthesis and degradation (Table 2). The two main genes identified in these metabolic pathways were phosphoribosylaminoimidazole carboxylase (PAICS) and purine nucleoside phosphorylase (PNP) which were both highly downregulated at all time points (Table S2). This confirms that combination dosing affects the biosynthesis of purine nucleotides. However, other metabolic pathways altered by combination dosing were not identified at gene level, suggesting the presence of a lag phase between gene-protein-metabolite expression or the insurgence of regulatory mechanisms directly operating at metabolite-protein level.

**Figure 6.**
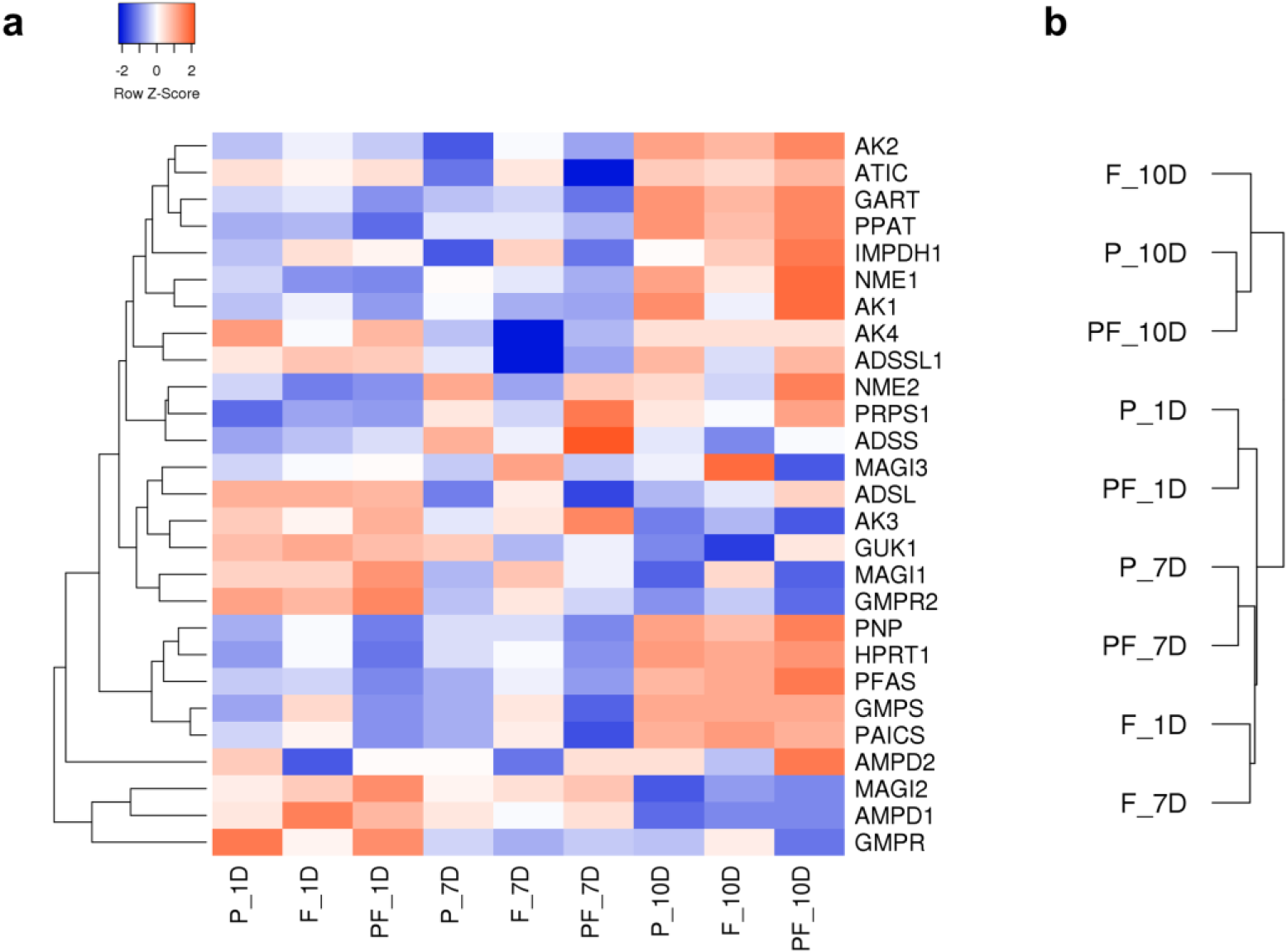
Differential regulation of selected genes after palbociclib (P), fulvestrant (F), and combination (PF) dosing, one day (1D), seven days (7D) and ten days (10D) postdose: a) heatmap of dysregulated genes (gene changes expressed as z-scores of the fold change), b) cluster analysis of gene expression profiles.

**Table 2.**
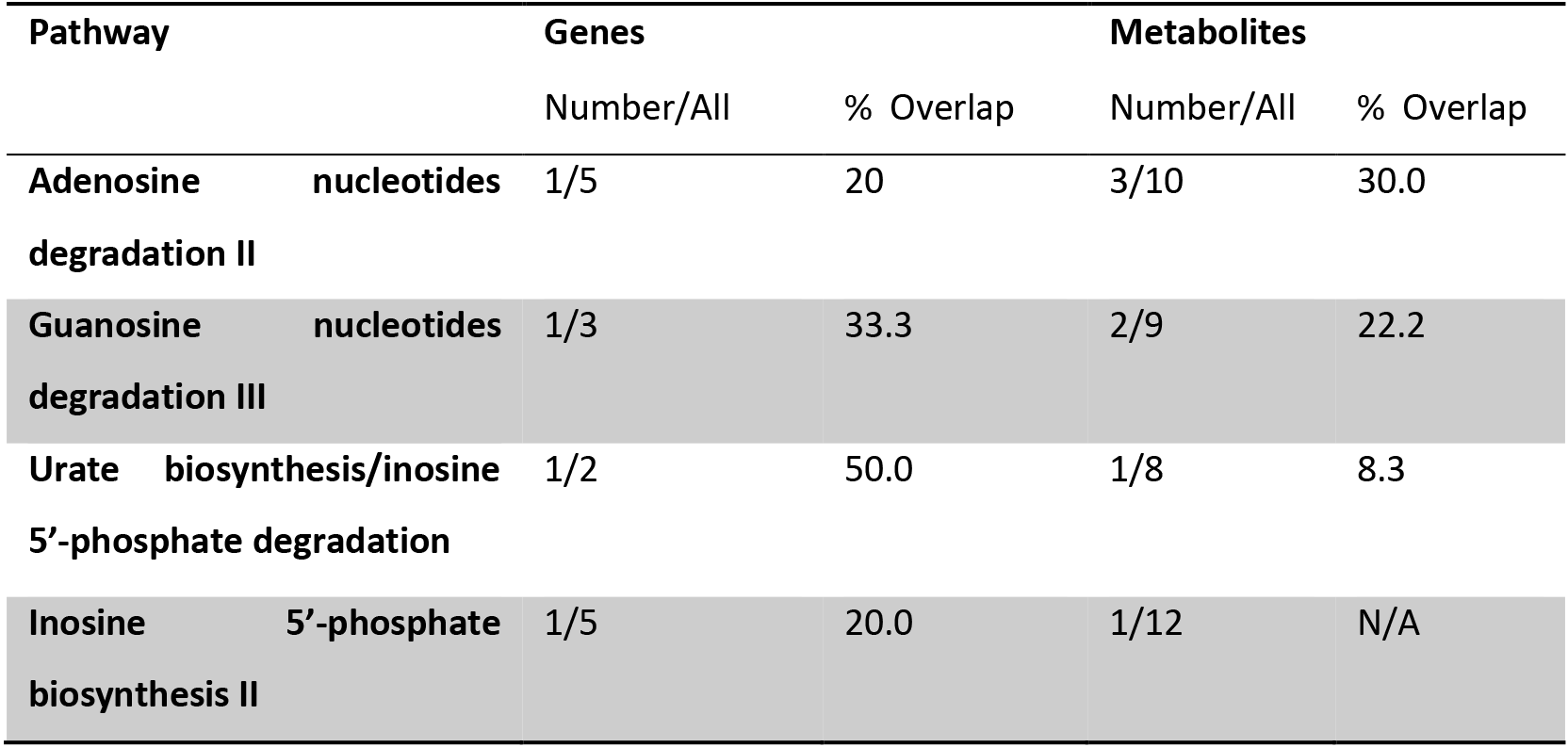
Active pathways mapped using untargeted metabolite and RNA-seq data at seven days post combination dose by XMCS Online. Pathway names provided by BioCyc Database Collection (Caspi et al., 2014).

### Comparison to previous results

We previously identified enhanced disruption to nucleotide metabolism, amino acids, and pentose phosphate pathway intermediates upon a 2-day combination therapy with palbociclib and letrozole. Similar to the single treatment with palbociclib, minor changes were seen to cell metabolism; slight decreases in malate, no changes to amino acids and the majority of central carbon metabolites. Single treatments with letrozole and fulvestrant appeared to have different effects on MCF-7 cells, underscoring their different endocrine mechanisms; fulvestrant is a selective ER inhibitor, whereas letrozole inhibits estrogen biosynthesis. The metabolites with the strongest response to fulvestrant after 2 days of dosing were sedoheptulose-7-phosphate, serine, 6-phosphogluconate, fumarate, malate, succinate, cholesterol sulfate and taurine. Whereas with letrozole, only slight metabolic responses were seen with changes to, 5-phosphogluconic acid, uridine, and metabolites in the glycolysis pathway.

Combination dosing with letrozole saw decreases in amino acids at 2 days, which were only apparent with fulvestrant at 7 days postdose. The major difference between the two treatments was the effect on nucleotide metabolism. In the case of fulvestrant combination dosing, we saw decreases in nucleotides, whereas with letrozole, increases were seen after 2 days of dosing, however it is not clear at this point whether a longer dosing time would result in similar effects. Thus, comparing these treatments show that combination therapies which include a CDK 4/6 inhibitor and endocrine therapy have a more profound effect on breast cancer cell metabolism.

## Conclusions

To advance the understanding of single versus combination drug therapeutics on cancer cell metabolism we used meta-analyses and a novel multi-omics technology to correlate metabolites and genes for deciphering dysregulated metabolic pathways. This tool freely available on XCMS Online revealed several pathways significantly modified by palbociclib, fulvestrant or a combination of the two. We observed that individually, palbociclib and fulvestrant caused disruption to shared metabolic pathways, however only fulvestrant acts on the PPP and has a more profound effect on amino acid biosynthesis. The combined effect of dosing with both drugs enables a comprehensive attenuation of metabolic pathways involved in coenzyme production, energy metabolism and macromolecule biosynthesis. This confirms the enhanced effect of combined breast cancer therapy over single treatment which was reported recently in a phase 3 study *in vivo* (Turner et al., 2015). Interestingly, after seven days post-treatment, we observed the initiation of a salvage mechanism, whereby the glycolytic pathway is shunted towards the PPP indicating that the cells may be attempting to synthesize serine and purines through an alternative mechanism. Thus, future studies should be designed to investigate the adaptive mechanisms of the cancer cells after removal of the drugs.

## Experimental procedures

### Cell culture

MCF-7 breast cancer cells (ATCC, Manassas, VA) were cultured in RPMI 1640 medium (Gibco-Life Tech, Grand Island, NY) supplemented with 10% fetal bovine serum (Sigma, St. Louis, MO) and penicillin-streptomycin (Gibco-Life Tech, Grand Island, NY) at 37°C and 5% CO2. Cells were passaged routinely at a ratio of 1:3 or 1:4 every 3-4 days using trypsin/EDTA. For metabolomics experiments, cells were seeded into 150 mm cell culture dishes (Corning, NY) and treated with either palbociclib (200 nM), fulvestrant (10 nM), a combination of those, or the vehicle as control. Four replicates were generated per experiment (approximately 3-5 million cells per replicate). Cells were harvested after two and seven days. Additionally, cells were taken after day seven with a refresh of the medium containing the respective drug(s) after day four. In total there were n=4/drug treatment group at two, seven, and seven days with refeeding, resulting in a total number of 48 biological samples for metabolomics analysis.

### Sample preparation for metabolomics experiments

Prior to sample harvest, cells were washed twice with phosphate buffer solution (Gibco-Life Tech, Grand Island, NY) and 500 μL water was added to the culture dish. The bottom of the culture dish was flash dipped into liquid nitrogen to quench metabolism immediately and cells harvested using a cell scraper. Cell suspensions were transferred to 1.5 mL Eppendorf tubes and further processed with three freeze-thaw cycles (1 min freezing, 5 min thawing) on wet ice using liquid nitrogen and 10 minutes of sonication in an ice water bath. Samples were centrifuged for 10 min at 13.000 rpm and 4°C. Supernatants were collected and protein concentration was quantified using a Pierce Micro BCA protein assay kit (Pierce, Rockford, IL). Lysed samples were further extracted in acetonitrile/methanol/lysate (2:2:1 v/v/v). Tubes were then vortexed for 30 s in 1.5 mL Eppendorf tubes, sonicated for 10 min and stored at −20°C for 1 h. Samples were subsequently centrifuged for 15 min at 13.000 rpm and 4°C. Supernatants were transferred to 1.5 mL high recovery glass autosampler vials (Agilent Technologies, Santa Clara, CA, United States) and dried in a speedvac (Labconco). According to the protein concentration, the samples were resuspended in acetonitrile/water (50/50 v/v), where the lowest protein concentration was resuspended in 100 μL, and all other samples were relatively adjusted thereafter. Samples were stored at −80°C until analysis.

### Untargeted metabolomics analysis

Analyses were performed using a high performance liquid chromatography (HPLC) system (1200 series, Agilent Technologies) coupled to an electrospray ionization (ESI) source and a 6550 ion funnel quadrupole time-of-flight (Q-TOF) mass spectrometer (Agilent Technologies). Samples were injected (8 μL) onto a Luna aminopropyl, 3 μm, 150 mm × 1.0 mm I.D. column (Phenomenex, Torrance, CA, United States) for HILIC analysis in ESI negative mode. HILIC was chosen to analyze predominantly central carbon metabolites as they are typically retained better by HILIC stationary phases upon comparison to reversed phase columns. Pooled samples were injected after three experimental samples whereas one solvent blank was injected after every sample for QC. Mobile phase was A = 20 mM ammonium acetate and 40 mM ammonium hydroxide in 95% water, 5 % acetonitrile and B = 95% acetonitrile, 5 % water. The linear gradient elution from 100 % B (0-5 min) to 100 % A (50-55 min) was applied in HILIC at a flow rate of 50 μL/min. To ensure column re-equilibration and maintain reproducibility a 10 min post-run was applied. ESI source conditions were set as follows: gas temperature 200°C, drying gas 11 L/min, nebulizer 15 psi, fragmentor 365 V, sheath gas temperature 300°C, sheath gas flow 9 L/min, nozzle voltage 500 V, capillary voltage 2500 V. The instrument was set to acquire over a *m/z* range from 60-1000 with the MS acquisition rate of 1.67 spectra/s. For the acquisition of MS/MS spectra of selected precursors the default isolation width was set to narrow (1.3 Da), with a MS acquisition rate at 1.67 spectra/s and MS/MS acquisition at 1.67 spectra/s. The collision energy was set to 20 eV. Data was processed using XCMS Online (Tautenhahn et al., 2012) with a p-value of < 0.05 and q-value of <0.1 set as statistical significance threshold cutoffs. Features were listed in a feature list table and as an interactive cloud plot, containing their integrated intensities (extracted ion chromatographic peak areas), observed fold changes across the sample groups, and statistical significance for each sample.

### Targeted metabolomics analysis

For targeted analysis, a volume of 2 μL was injected onto a Luna aminopropyl, 3 μm, 150 mm × 2.0 mm I.D. column (Phenomenex) using an Agilent Technologies series 1290 Infinity HPLC system with a gradient mobile phase of A = 20 mM ammonium acetate and 40 mM ammonium hydroxide in 95% water, 5% acetonitrile and B = 95% acetonitrile, 5 % water. The linear gradient elution from 95% B (0-2 min) to 10% B for 13 min, then to 0% B in 2 min, held at 0% B for 3 min, and equilibrated back to 95 % B over 4 min at a flow rate of 350 μL/min. Quantification of metabolites was performed by dynamic multiple reaction monitoring triple quadrupole mass spectrometry (Agilent Ion-Funnel 6490). The ESI source conditions were as following: gas temperature 225°C, gas flow 15 L/min, nebulizer 35 psi, sheath gas 400°C, sheath gas flow 12 L/min, capillary voltage 2500 V (ESI negative) or 3500 V (ESI positive) and nozzle voltage 0 V. The cycle time was set to 500 ms. The collision energies, quantifier and qualifier ion transitions were optimized for each metabolite using MassHunter Optimizer software and are reported in the supplementary Table S1. To ensure accurate quantification, external calibration with standard compound mixtures was performed. Agilent QQQ Quantitative Analysis software was used to calculate the absolute concentrations of the metabolites in the samples.

### Transcriptome analysis

MCF-7 breast cancer cells were grown as above and collected at various time points (one day and seven days after dosing, with an additional time point at ten days with re-dosing at day seven), n=3/group. Whole transcriptome RNA sequencing was performed by Biomiga (San Diego, CA). Each treatment and time point combination was profiled in biological duplicates. The 50bp paired end reads were mapped by bowtie2 (Langmead and Salzberg, 2012) and quantified using RSEM package (Li and Dewey, 2011). Differential expression statistics was determined with EdgeR algorithm (Robinson et al., 2010) from expected counts. Gene set enrichment analysis (Subramanian et al., 2005) was performed using TPM values based on weighted signal-to-noise metric and false discovery rate (FDR) was assessed from 1000 permutations. Gene expression profiles cluster analysis was performed by average linkage with the Euclidean distance measurement method. Pathway gene signatures were prepared by mapping gene pathway list from KEGG database (Kanehisa et al., 2012; Ogata et al., 1999) to Entrez gene IDs at Pfizer.

### Stats

GraphPad Prism v 6.00 (GraphPad Software, Inc, San Diego, CA) was used for statistical analysis. The quantitative triple quadrupole data was log transformed and expressed as mean ± standard error of the mean (S.E.M) after two-tailed t-tests were carried out. Comparisons with p < 0.05 were assigned to be statistically significant and noted on each graph.

## Author contributions

CHJ, BW and LH extracted, analyzed and validated the metabolomics data. AP contributed in data analysis and interpretation. NVL carried out the cell culture and drug dosing. ZZ created and analyzed the transcriptomics data. All the authors gave input into experimental design, data analysis and writing the paper.

## Funding

The authors further thank and the Austrian Science Fund (FWF; Erwin Schrödinger Fellowship J-3808 awarded to B.W.) for financial support.

## Conflict of Interest

Nathan V Lee, Zhou Zhu, Anthony Mazurek, Stephen Dann and David Shields are employees of Pfizer. Valeria Fantin is a current employee of Oric Pharmaceuticals.

